# Label-Free Quantification of Apoptosis and Necrosis Using Stimulated Raman Scattering Microscopy

**DOI:** 10.1101/2025.03.01.641010

**Authors:** Shivam Mahapatra, Shreya B. Shivpuje, Helen C. Campbell, Boyong Wan, Justin Lomont, B. Dong, Seohee Ma, Karsten J. Mohn, Chi Zhang

**Affiliations:** James Tarpo Jr. and Margaret Tarpo Department of Chemistry, Purdue University, 560 Oval Dr., West Lafayette, IN 47907, USA; Department of Statistics, Purdue University, 150 N University St, West Lafayette, IN 47907, USA; Merck & Co. Inc; Purdue Institute for Cancer Research, 201 S. University St., West Lafayette, IN 47907, USA; Purdue Institute of Inflammation, Immunology, and Infectious Disease, 207 S. Martin Jischke Dr., West Lafayette, IN 47907, USA

**Author notes:** Sh.M. and S.S., these authors contribute equally to this work.

## Abstract

Recombinant proteins are critical for modern therapeutics and diagnostics, with Chinese hamster ovary (CHO) cells serving as the primary production platform. However, environmental and chemical stressors in bioreactors often trigger cell death, particularly apoptosis, posing a significant challenge to recombinant protein manufacturing. Rapid, label-free methods to monitor cell death are essential for ensuring better production quality. Stimulated Raman scattering (SRS) microscopy offers a powerful, label-free approach to measure lipid and protein compositions in live cells. We demonstrate that SRS microscopy enables rapid and reagent-free analysis of apoptotic and necrotic transitions. Our results show that apoptotic cells exhibit higher protein concentrations, while necrotic cells show an opposite trend. To enhance analysis, we developed a quantitative single-cell analysis pipeline that extracts chemotypic and phenotypic signatures of apoptosis and necrosis, enabling the identification of subpopulations with varied responses to stressors or treatments. Furthermore, the cell death analysis was successfully generalized to other stressors and cell types. This study highlights SRS microscopy as a robust and non-invasive tool for rapid monitoring of live cell apoptotic and necrotic transitions. Our method and findings hold potential for improving quality control in CHO cell-based biopharmaceutical production and for evaluating cell death in diverse biological contexts.

## Introduction

Recombinant proteins, such as monoclonal antibodies (mAbs), cytokines, hormones, and enzymes, provide better efficacy and higher stability for therapeutic and diagnostic purposes and therefore are key products in the modern pharmaceutical industry. Chinese hamster ovary (CHO) cells are the workhorse of the current pharmaceutical industry in producing these recombinant proteins.^1–2^ These CHO cells are typically grown in bioreactors for mass production. The yield of recombinant proteins is highly influenced by the healthy condition of cells. Apoptosis, which can be triggered due to cell aging or environmental stimuli, can significantly impact CHO cell viability and protein production.^3–4^

In a healthy organism, apoptosis is a well-regulated self-destructive mechanism of cells for maintaining overall homeostasis by removing damaged, abnormal, aged, or unnecessary cells.^5–6^ However, apoptosis can also be triggered by external stimuli.^7–8^ Such external impact poses a key challenge for CHO cells in the production of recombinant proteins.^3^ For example, oxygen imbalance, temperature fluctuation, pH change, nutrition depletion, shear stress, etc. can cause CHO cell apoptosis in bioreactors.^3,9^ Uncontrolled apoptosis during antibody production can significantly reduce protein yields and contaminate the final antibody product with remnants of dead host cells. Being able to evaluate the health condition of CHO cells inline or online and detect apoptosis at an early stage during bioreactor production is essential to ensure high yields of recombinant proteins. Furthermore, identifying the cell clones that are more resistant to apoptosis would help large-scale manufacturing.

Apoptosis proceeds via a series of characteristic changes such as cell shrinkage, chromatin condensation and DNA fragmentation, enrichment of proteins in a small volume, and finally, membrane blebbing, followed by release of apoptotic bodies.^6^ If apoptosis is prolonged, it may result in necrotic changes known as secondary necrosis.^10–11^ Investigating these changes, both molecular and morphological, allows researchers to understand how cells respond to different stressors. Conventionally, molecular changes can be monitored using caspase activity assays^12^, DNA fragmentation assays^13^, Annexin V or Propidium Iodide staining^14–15^, western blotting^16^, and immunostaining^17^. Although these assays are highly sensitive to apoptosis and provide early-stage information about the pathways involved, they might process time ranging from hours to days. This long process and turnover time pose a strong challenge for rapid analysis and decision-making, which are extremely valuable for making fast adjustments or decisions for protein production. Moreover, some methods might require fixing samples, compromising the information of live cells, and adding processing time. Probing morphological changes, on the other hand, does not require any specialized reagents and can enable faster measurements.^18^ However, it lacks chemical information to verify the apoptotic transition or evaluate the potential causes of the changes. There is a critical need to develop a rapid, label-free, yet chemically selective method to analyze CHO cell death. Chemical imaging by stimulated Raman scattering (SRS) microscopy provides a unique way to fulfill this requirement.

SRS microscopy utilized Raman signals to quantify protein and lipid compositions in cells.^19–25^ The imaging speed can be as fast as fluorescence microscopy. The label-free imaging scheme obviates the need for labeling or pre- and post-processing of the samples. Using SRS microscopy, we identified an increase in intracellular protein concentration as a critical marker for evaluating apoptotic changes in CHO cells. Detection of apoptosis, in both adherent and suspended cells, can be achieved within 30 minutes. Additionally, we observed that necrotic transitions are characterized by a decrease in intracellular protein concentration, which can be distinguished from apoptotic and healthy cells. We evaluated various apoptotic and necrotic inducers, including staurosporine, hydrogen peroxide, dimethyl sulfoxide (DMSO), and blue light. To enhance analysis, we developed a single-cell analysis pipeline integrating deep-learning and CellProfiler. This pipeline extracts key chemotypic and phenotypic signatures of cell death at the single-cell level, revealing unique characteristics and features associated with apoptotic and necrotic transitions that are not detectable through ensemble cell analysis. This work establishes SRS microscopy as a powerful and rapid method for evaluating cell death. The methodology and findings hold significant promise for enhancing quality control in biopharmaceutical production and providing new insights into cellular responses to various stimuli.

### Experimental Section

#### Stimulated Raman scattering microscopy

The SRS microscope used in this study has been described previously.^26–27^ Briefly, a femtosecond dual-output laser (InSight X3+) was employed, with one output fixed at 1045 nm and the other tunable between 690–1200 nm for SRS signal excitation. For imaging cells in the C-H vibrational range, the tunable output was set to 802 nm, corresponding to a Raman shift centered at 2900 cm^-1^. The laser pulses were chirped to picoseconds using glass rods before entering the microscope. The Stokes laser at 1045 nm was modulated at 2.4 MHz using an acousto-optic modulator. SRS signals were detected with a silicon photodiode (S3994, Hamamatsu) and demodulated using a lock-in amplifier (HF2LI, 50 MHz, Zurich Instruments). A 2D galvo scanner enabled image acquisition at 10 µs per pixel, resulting in a 400×400-pixel frame captured in approximately 2 seconds. Samples were mounted on a 3D motorized stage, which allowed the stitching of single-frame images into larger areas. Motorized axial adjustment facilitated 3D imaging by sweeping the objective lens’s axial position. A 40x water immersion objective lens (LUMPLFLN 40XW, NA=0.8, Olympus) was used for imaging. Image acquisition and display were managed by lab-written LabVIEW-based software.

For the blue laser treatment of cells, a 100 mW, 405 nm continuous-wave (CW) laser was collinearly combined with the infrared femtosecond laser pulses and scanned within the same field of view using the 2D galvo scanner. The laser power at the sample for treatment was either 2 mW or 1 mW.

#### Quantitative cell analysis (in brief for manuscript)

The obtained SRS images were preprocessed using ImageJ to facilitate ensemble cell analysis or prepare for single-cell analysis. For each treatment condition, 25 images (80 µm × 80 µm) were acquired and processed, with the total number of cells per condition ranging from 140 to 700. The 2D and 3D SRS images presented in the figures were processed using ImageJ. To better visualize intensity changes under various conditions, most SRS images were displayed using a pseudo-color Jet color scheme.

LDs were identified from C-H SRS images by subtracting a Gaussian-blurred version of the original image (radius = 2 pixels) from the original image.^28^ The LD masks generated through this process could be multiplied with the original C-H SRS image to isolate the LD contents. Alternatively, LDs could be removed from the C-H SRS images by subtracting the LD contents from the original image. The original C-H SRS images were used as input for Cellpose, enabling automated and supervised cell segmentation with high accuracy. The cell masks generated by Cellpose were combined with the SRS images and served as input for CellProfiler.

In CellProfiler, a custom pipeline was developed to extract features associated with individual cells. The features analyzed in this study included cell size, eccentricity, average SRS intensity, average protein intensity, integrated SRS intensity, and integrated LD intensity. Histograms of these features were plotted and normalized for each condition using ImageJ and Origin for comparative analysis.

Additionally, 2D contour plots correlating different parameters, such as cell size and average SRS intensity, were generated using custom MATLAB scripts. For LD mobility analysis, 50 frames of time-lapse SRS images were acquired. The LD trajectory quantification methods followed those described in a previous publication.^29^.

The details of the analysis methods can be found in the supporting information.

#### Cell preparation

CHO-K1 cells were purchased from ATCC for the apoptosis studies. Cells were cultured in F-12K Medium (Kaighn’s Modification of Ham’s F-12 Medium) with 10% fetal bovine serum (FBS, ATCC) and 0.5% penicillin/streptomycin (Thermofisher Scientific). The cells were plated in small glass-bottom dishes (MatTek Life Sciences) with 2 mL culture medium and then incubated in a CO_2_ incubator at 37 °C and 5% CO_2_ concentration. The cells were allowed to grow to approximately 50-70% confluency before using for SRS microscopy in live cells. HeLa cells were obtained from ATCC for apoptosis studies in mammalian cells. The cells were cultured in Dulbecco’s Modified Eagle Medium (DMEM, ATCC) with 10% fetal bovine serum (FBS, ATCC) and 1% penicillin/streptomycin (ThermoFisher Scientific) to reach approximately 50-70% confluency before imaging. HeLa cells transfected with EB3-EGFP were obtained from Biohippo and were cultured similarly to normal HeLa cells.

#### Chemical treatment of cells

CHO-K1 or HeLa cells are first seeded in respective glass-bottom dishes and allowed to grow to a confluency level of about 50-70%. Staurosporine was added to the culture medium at a final concentration of 1 µM and the cultures were incubated at 37 °C and 5% CO_2_ concentration before imaging. The cells were imaged alive. For H_2_O_2_ treatment, a final concentration of 5 mM of H_2_O_2_ was added to the culture medium. In both cases, the incubation duration was varied to evaluate the correlation between the extent of apoptosis and incubation duration. In the H_2_O_2_-treatment case, the media was removed, and the cells were washed twice with phosphate-buffered saline (PBS) and fixed with 10% formalin before imaging.

For suspended cell treatment, CHO-K1 cells were suspended by trypsin and treated with 1 µM staurosporine at different time lengths in centrifuge tubes in the CO_2_ incubator.

## Results

### SRS detection of apoptosis progression in adherent CHO cells induced by staurosporine

CHO-K1 cells were chosen for the majority of this research. To induce apoptosis, cells were incubated with 1 µM staurosporine, a widely used apoptosis inducer^30^, for varying durations. Staurosporine induces apoptosis majorly by activating the caspase cascade, inducing mitochondria damage, and inhibiting protein kinases. SRS microscopy was conducted by measuring the C-H stretching region centered at 2910 cm^-1^, which revealed both lipid droplets (LDs) and intracellular proteins. To minimize intensity variations, the same treatment group, along with the control group, was imaged consecutively on the same day with the same microscope and laser conditions. The laser focus was maintained at a plane just above the glass substrate inside the cells.

As shown in **Figures 1A and 1B**, the SRS intensity from adherent CHO cells increased steadily with prolonged staurosporine treatment. The SRS signals captured in these images consist of contributions from both intracellular LDs, visible as small, bright, high-intensity dots, and overall cellular proteins, which dominate the remaining cellular regions. Nucleic acid signals were negligible compared to those from LDs and proteins at this wavenumber. To distinguish LDs from proteins, a quantitative image analysis method was employed to exclude LDs from the overall cell images, as demonstrated in **Figures 1C and 1D** and detailed in the supporting information. Images at different time points were analyzed to compare the average SRS signals from cells in each image. This analysis revealed a statistically significant increase in SRS intensity as a function of treatment time. When LD signals were excluded, leaving only protein signals for analysis, a similar trend was observed. These findings indicate that intracellular protein concentration significantly increases in CHO cells undergoing staurosporine-induced apoptosis.^24^ A similar increase in protein concentration during staurosporine-induced apoptosis is also observed in other cell types, such as HeLa cells (**Figure S1**).

**Figure 1.**
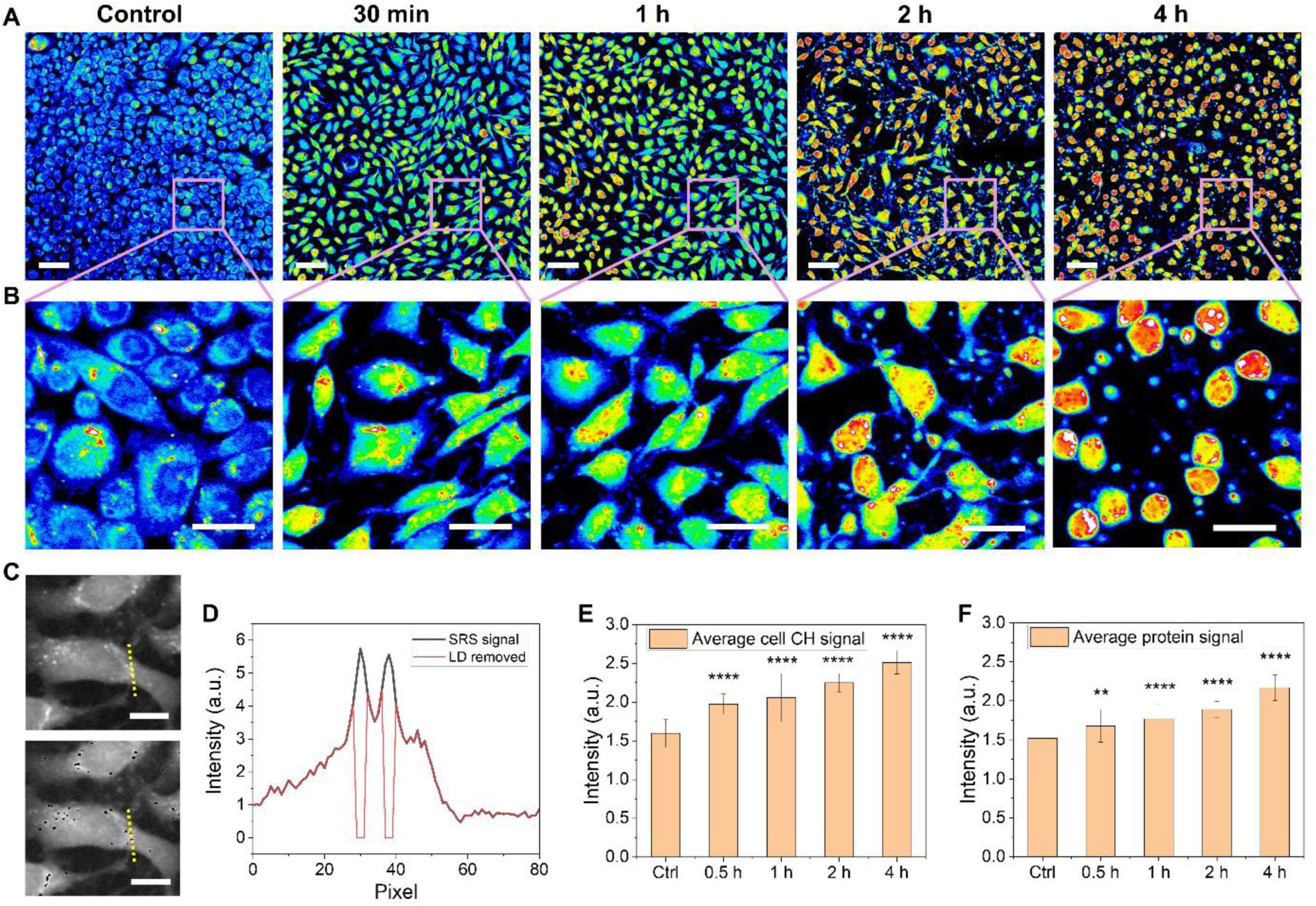
SRS detects increased intracellular protein concentration during staurosporine-induced apoptosis of adherent CHO cells. (A) SRS images of adherent CHO-K1 cells under untreated (control) and 1 µM staurosporine-treated conditions, captured at time points ranging from 30 minutes to 4 hours. Images are displayed using a jet color scheme. (B) Magnified views of selected regions from panel A. (C) SRS images presented in grayscale (top) and the corresponding images after lipid droplet removal. (D) SRS intensity profiles plotted along the dotted lines shown in panel C. (E) Average cellular SRS intensity quantified from images in panel A under various conditions. (F) Average cellular SRS intensity calculated after lipid droplet removal from the images. Scale bars: 50 µm in panel A, 20 µm in panel B, and 10 µm in panel C. ** p<0.01, **** p<0.0001

### Statistical analysis of the phenotypic and chemotypic changes at the single-cell level

While the ensemble cell analysis in **Figure 1** highlights the trend of increasing protein concentration during apoptosis, it lacks single-cell information. This approach does not segment cells and provide detailed information on the phenotypic and chemotypic traits of individual cells. To overcome these limitations, we developed a single-cell analysis pipeline utilizing Cellpose^31^ and CellProfiler^32^ for automated segmentation of individual cells and statistical extraction of diverse physical and chemical features. Cellpose is an advanced deep-learning-assisted cell segmentation tool, while CellProfiler is a robust software widely used for single-cell analysis and has been successfully applied with SRS microscopy.^25^ The workflow of the new statistical analysis pipeline we developed in this work is detailed in **Figure 2A**. Cellpose 2.0 was used to segment cells from raw SRS images.^33–34^ Additionally, the SRS images were processed to separate LDs from cellular proteins using a previously reported method.^28^ These processed images—raw SRS, LD-specific, or LD-removed—along with Cellpose-generated cell masks, served as inputs for CellProfiler. A pipeline was established to extract features such as cell size, eccentricity, total SRS intensity, average SRS intensity, LD content, and more, enabling the statistical analysis of single-cell data for various treatment conditions. Histograms of cell counts based on SRS intensity, cell size, eccentricity, etc., can be generated and plotted using Origin, while MATLAB was used to create 2D density contour plots correlating different parameters.

**Figure 2.**
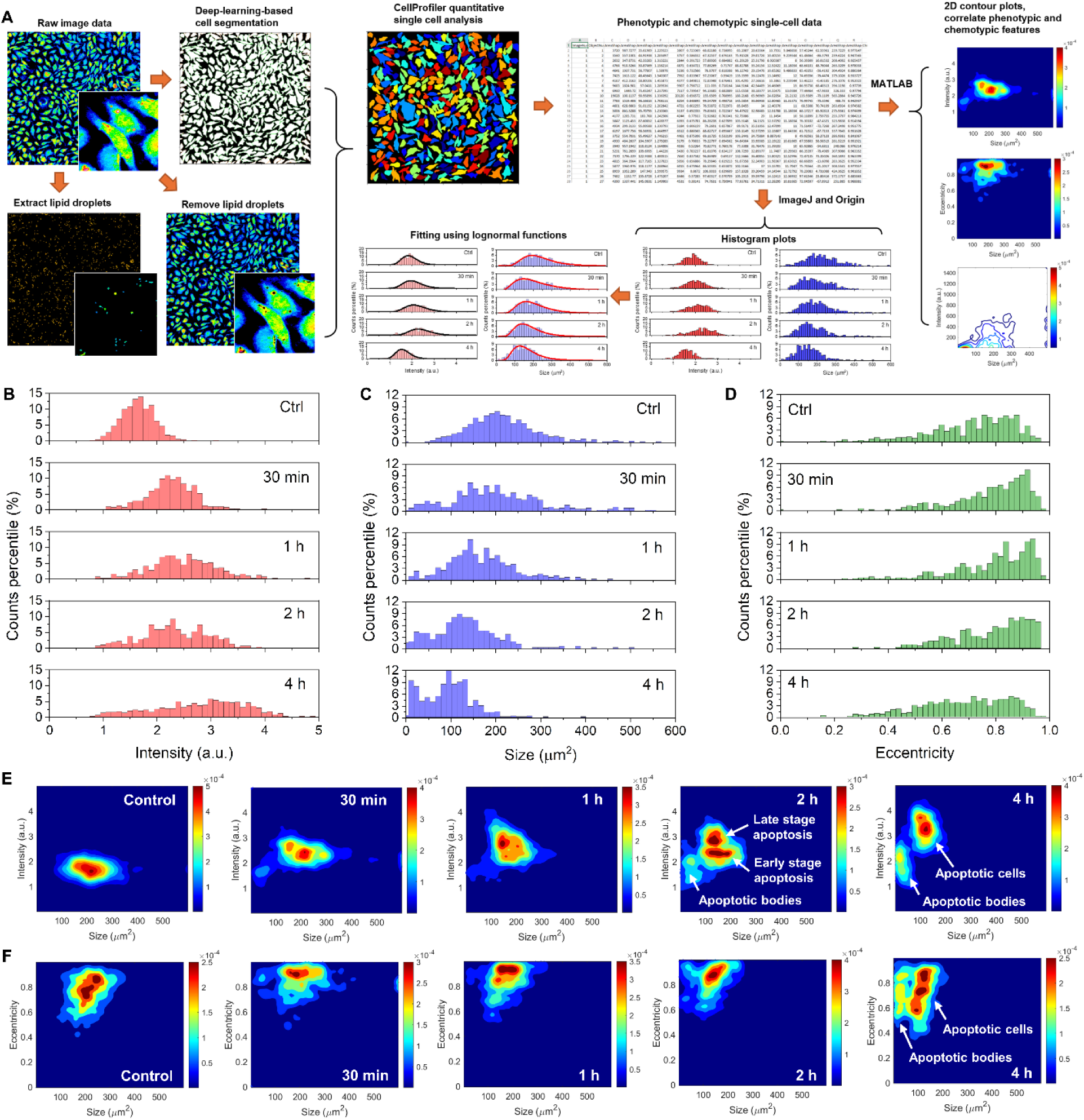
Single-cell quantitative analysis of CHO-K1 cell apoptosis transitions induced by 1 µM staurosporine. (A) Schematic workflow illustrating the single-cell quantitative analysis method developed and employed in this study. (B) Histograms showing the distribution of cellular average SRS intensity. (C) Histograms showing the distribution of cell sizes. (D) Histograms showing the distribution of cell eccentricity. (E) Two-dimensional contour plots depicting the relationship between average SRS intensity and cell size under various apoptotic conditions. The plots are presented using a jet color scheme, with distinct object populations labeled in the 2-hour and 4-hour time points. (F) Two-dimensional contour plots showing the relationship between cell eccentricity and cell size across different apoptotic conditions.

Using this single-cell analysis pipeline, we re-examined adherent CHO cells treated with staurosporine (**Figure 1**). Quantitative results revealed a similar trend in the increase of average SRS intensity as a function of treatment time (**Figure 2B**). Additionally, the intensity distribution broadened over time, suggesting heterogeneity in the cellular response. When LDs were excluded, the analysis still showed a consistent increase in SRS signal during apoptosis (**Figure S2**), indicating that the overall intracellular protein concentration increases in apoptotic CHO cells, with minimal influence from LDs on whole-cell intensity measurements. Further analysis quantified the integrated total cell SRS intensity (**Figure S3**), showing a slight increase during the early stages of apoptosis. This increase is caused by the rise in protein concentration and changes in cell volume in 3D. A detailed explanation can be found in the supporting information. After 4 hours of treatment, the intensity returned to control levels, indicating the loss of cellular proteins at this time point. These findings suggest that apoptosis does not significantly reduce cellular protein content until 4 hours of staurosporine treatment.

Beyond changes in intensity, we observed that CHO cells tend to decrease in size during apoptosis, as shown in **Figure 2C**. This response is generally known as apoptotic volume decrease (AVD).^35–36^ A new population of objects smaller than 1500 pixels (∼60 μm²) emerged. This size is too small for a single CHO cell, suggesting that these particles correspond to apoptotic bodies^37^, which are visible in **Figure 1B**. Our results demonstrate that the number of apoptotic bodies increases with staurosporine treatment time. Additionally, we quantified cell eccentricity, where higher values (approaching 1) indicate elongation and lower values reflect a more circular shape. As shown in **Figure 2D**, staurosporine-induced apoptosis increased cell polarity between 30 minutes and 2 hours of treatment, indicating early-stage apoptosis. By 4 hours, the cells reverted to a more circular shape, indicating late-stage apoptosis. These changes in cell size and shape, indicative of different apoptotic stages, cannot be captured through ensemble analysis of SRS images.

To better correlate the physical and chemical features of single cells, we generated 2D density contour plots of cell size versus average SRS intensity under different treatment conditions, as shown in **Figure 2E**. The control group exhibits a single population with a size centered around 210 µm^2^ and an intensity centered at 1.8 (a.u.). Staurosporine treatment induces a progressive decrease in cell size accompanied by an increase in cell intensity over time. The emergence and growth of apoptotic bodies are distinguishable in these contour plots. Furthermore, late-stage apoptotic cells (higher intensity and smaller size) are well-differentiated from early-stage apoptotic cells, particularly at the 2-hour time point. These findings explain the broad SRS intensity distribution observed in **Figure 2B** during apoptosis, which results from the simultaneous presence of apoptotic bodies and cells at various apoptotic stages.

In the eccentricity versus cell size plots, two distinct populations of cells with different shapes were observed in the control group, with similar proportions in each. Staurosporine-induced apoptosis significantly increased cell eccentricity while reducing cell size, leading to a predominance of elongated cells between the 30-minute and 2-hour time points. By 4 hours, a substantial number of apoptotic cells had become smaller and more circular. These smaller, rounded cells, identified in the eccentricity versus intensity plots (**Figure S4**), correspond to more late-stage apoptotic cells.

Additionally, using LD-specific SRS images derived from raw SRS data, we quantified changes in LD content during apoptosis. A subset of cells exhibited increased LD counts at the 4-hour mark (**Figure S5**). Collectively, these single-cell analysis results highlight phenotypic and chemotypic changes during apoptosis, particularly distinguishing these changes across different subpopulations of cells.

### Apoptosis induced by other stressors

In addition to staurosporine, hydrogen peroxide (H_2_O_2_) is another commonly used inducer of apoptosis which induces apoptosis by oxidative stress and damage of membrane structures.^38–39^ CHO cells were treated with 5 mM H_2_O_2_ and analyzed at 0.5, 1, 2, and 4-hour time points. Single-cell analysis revealed similar trends of increasing average SRS intensity up to 2 hours, followed by a decrease at the 4-hour time point (**Figures 3A, B**). Cell size showed a continuous decline over time (**Figure 3C**). To better compare the changes in histograms, we fit the data with lognormal functions. The details of the fitting function and key parameters are detailed in the supporting information. The medium value can be used to compare the changes in intensity and cell size of various conditions. Correlated intensity versus cell size contour plots are presented in **Figure 3D**.

**Figure 3.**
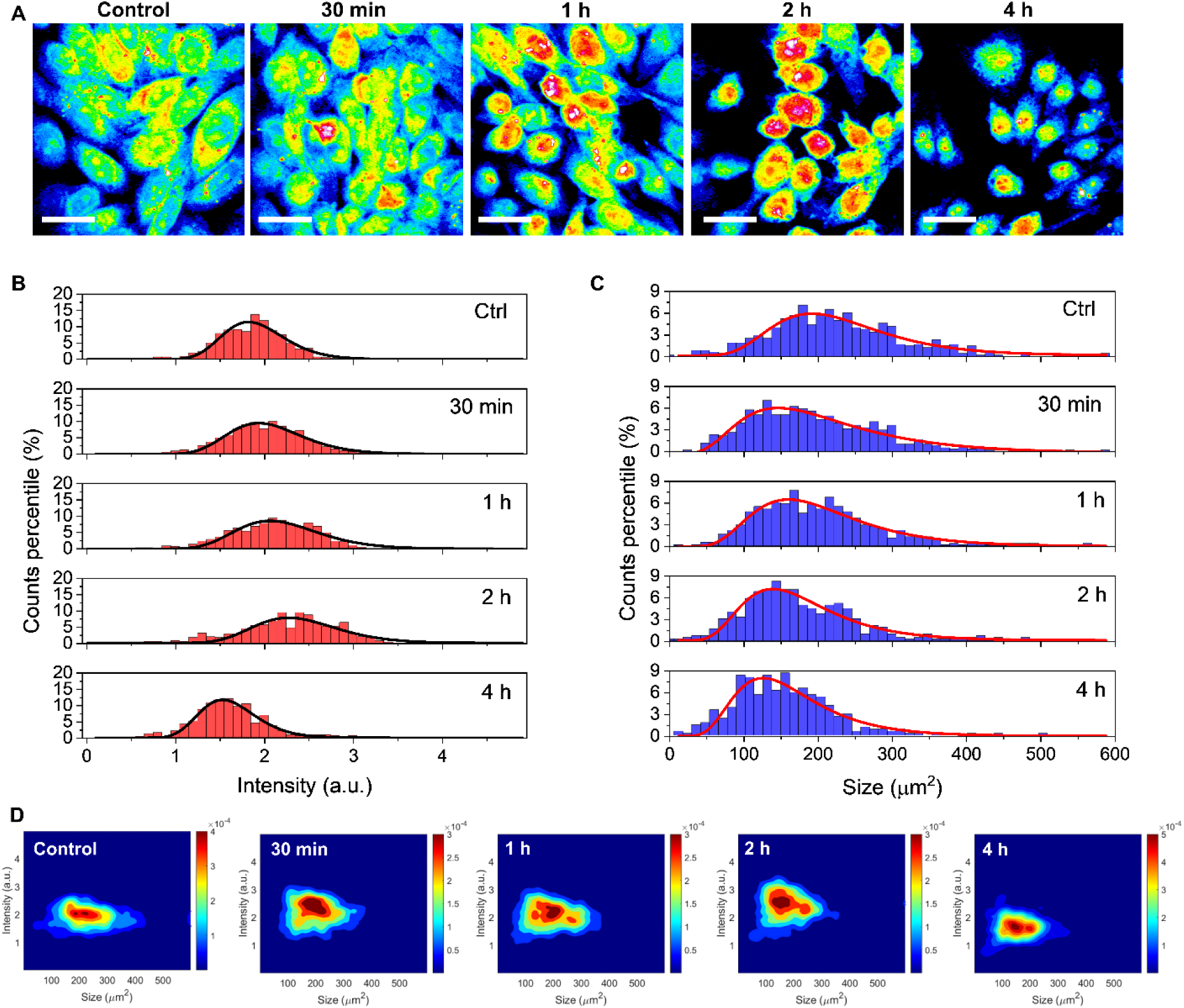
Cellular features during hydrogen peroxide-induced apoptosis and necrosis of adherent CHO cells. (A) SRS images of CHO-K1 cells under untreated (control) and 5 mM H_2_O_2_-treated conditions, captured at time points ranging from 30 minutes to 4 hours. Images are presented in a jet color scheme. (B) Histograms depicting the distribution of cellular average SRS intensity, with lognormal function fits. (C) Histograms showing the distribution of cell sizes, also fitted with lognormal functions. (D) Two-dimensional contour plots illustrating the relationship between average SRS intensity and cell size at various time points. The plots are displayed in a jet color scheme. Scale bar: 20 µm in panel A.

Compared to staurosporine-induced apoptosis, H_2_O_2_-induced apoptosis also demonstrated an increase in cellular protein concentration. The significantly fewer apoptotic bodies observed at the 2-hour time point can be attributed to an additional washing process performed before imaging. Moreover, the rate of protein concentration increase was less pronounced than in staurosporine-treated cells. These findings suggest that, while both stressors share a common trend of protein concentration increase during apoptosis, CHO cells respond to each inducer in distinct ways. Notably, the reduction in protein concentration at the 4-hour mark is unique to H_2_O_2_ treatment and contrasts with the staurosporine case. This decline in SRS signal is likely associated with secondary necrosis following prolonged H_2_O_2_ exposure, a phenomenon reported to occur in various cell types subjected to H_2_O_2_ treatment.^39–41^ We believe these cells are undergoing necrosis, as evidenced by the presence of empty vacuoles appearing as dark droplets and large membrane blebs, which are characteristic features of necrotic cells (**Figure S6**).^42–43^

### Differentiation apoptosis and necrosis using SRS microscopy

To confirm that reduced average SRS intensity is associated with necrosis, we induced necrosis by applying a high concentration of DMSO. A drop of DMSO was added to the culture dish, followed by rapid shaking to ensure homogeneous mixing with the culture medium. Although the final DMSO concentration was only 0.1%, the initial high concentration at the drop-cast area induced necrosis within 2 seconds of exposure. This rapid effect occurs because high DMSO concentrations irreversibly damage cell membranes.

We observed that this drop-casting and mixing method resulted in necrosis at the center of the drop-cast area, characterized by a slightly lower average SRS intensity (**Figures 4A, B**). In this region, intracellular vacuoles and significant swelling of necrotic cell membranes are evident. Surrounding the necrotic core was a belt of mixed necrotic and apoptotic cells, with apoptotic cells displaying high average SRS intensity (**Figures 4A, B**). Beyond this boundary, unaffected cells exhibited features similar to the untreated control group (**Figures 4A, B**). For comparison, **Figures 4C,D** present SRS images of control and 1-hour 1 µM staurosporine-treated CHO cells acquired on the same day.

**Figure 4.**
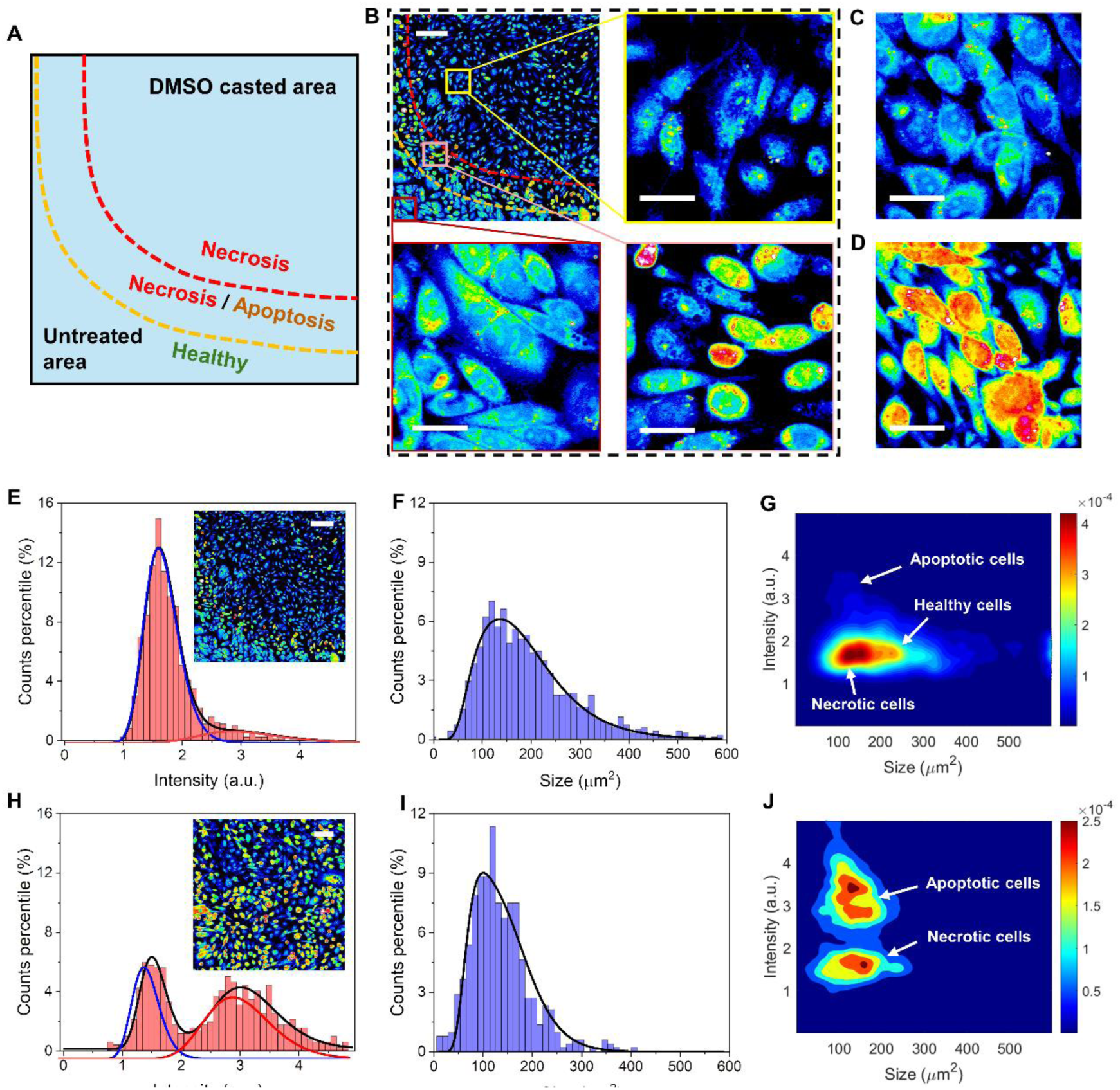
Distinguishing necrotic cells from apoptotic cells. (A) Schematic illustrating different regions of a culture dish following high-concentration DMSO exposure and rapid mixing. (B) SRS images of CHO-K1 cells showing necrotic, apoptotic, and healthy states at different areas of the dish after DMSO treatment. (C) SRS images of healthy CHO-K1 cells acquired on the same day as panel A. (D) SRS images of CHO-K1 cells treated with 1 µM staurosporine for 1 hour, also acquired on the same day as panel A. (E) Intensity histogram generated from single-cell analysis of the SRS image in panel B. Two distinct cell populations corresponding to apoptotic and other cells are identified in the intensity domain. The histogram is fitted with a dual-lognormal function. (F) Histogram of cell sizes derived from the SRS image in panel B, fitted with a tri-lognormal function. (G) Two-dimensional contour plots showing the relationship between average SRS intensity and cell size for the cells in panel B. (H-J) Similar analyses to panels E-G, performed on SRS images containing only apoptotic and necrotic cells. These populations are clearly separable based on average SRS intensity. Scale bars, 100 µm in panel B top left and panel E, 20 µm in the rest of panel B, C, and D, 50 µm in panel H. In panels E, F, H, I, black curves are fitting results, blue, red, and green curves are single-lognormal functions.

Statistical single-cell analysis revealed a broader intensity distribution. A subpopulation with higher intensity values corresponds to apoptotic cells within the population (**Figure 4E**). Although the separation between necrotic and healthy cells was minimal, these groups together were distinct from the apoptotic population (**Figure 4E**). To analyze the data, we applied a dual-lognormal function to fit the intensity distribution, estimating that approximately 12% of the cells in **Figure 4B** were apoptotic. The broad size distribution in the cell size histogram reflects the simultaneous presence of necrotic, apoptotic, and healthy cells. These three populations can be identified through lognormal fitting of the cell size histogram (**Figure 4E**). Using previously established median values for healthy and apoptotic cells, we estimated the ratio of cells in each population to be 16:83:20 (apoptotic:necrotic:healthy). Additionally, the 2D density contour plots in **Figure 4G** more effectively separated these populations. Notably, the necrotic cell population shifted toward smaller size values compared to healthy cells, with only a slight decrease in median intensity.

To further validate our quantitative analysis, we performed a similar experiment by drop-casting DMSO containing staurosporine onto CHO cells, followed by rapid mixing to achieve a final staurosporine concentration of 1 µM for 1 hour of incubation. We identified regions containing a mixture of necrotic and apoptotic cells, with no healthy cells present. In this scenario, necrotic and apoptotic cells were well-separated in the intensity domain (**Figure 4H**), with a rough population ratio of 3:5. While the cell size distributions of necrotic and apoptotic cells overlapped, they were distinguishable using lognormal distribution fitting (**Figure 4I**). The 2D contour plots in **Figure 4J** showed a clear separation between necrotic and apoptotic cells, with distinct early- and late-stage apoptotic subpopulations.

Another method to induce cell death is through intense blue light irradiation, which generates reactive oxygen species (ROS) inside cells through interactions with intrinsic biomolecules such as nicotinamide adenine dinucleotide hydrogen (NADH), flavin adenine dinucleotide (FAD), and cytochromes. **Figure 5A** illustrates the approach used to induce necrosis with a 10 mW 405 nm laser, followed by SRS imaging of a larger area encompassing the treated region. The treatment was performed continuously for 50 frames (80 s in total) with a pixel dwell time of 10 microseconds. Comparing SRS images taken before (**Figure 5B**) and after (**Figure 5C**) blue light irradiation, we observed a slight decrease in SRS intensity in CHO cells. To further investigate these radiated cells, we performed 3D imaging of the blue laser-treated regions, revealing cell membrane swelling and the formation of large membrane blebs, both hallmark features of necrotic transitions. Such necrosis is likely induced by the excessive ROS generation inside cells that damage cellular organelles, DNA, and plasma membranes.^44–47^

**Figure 5.**
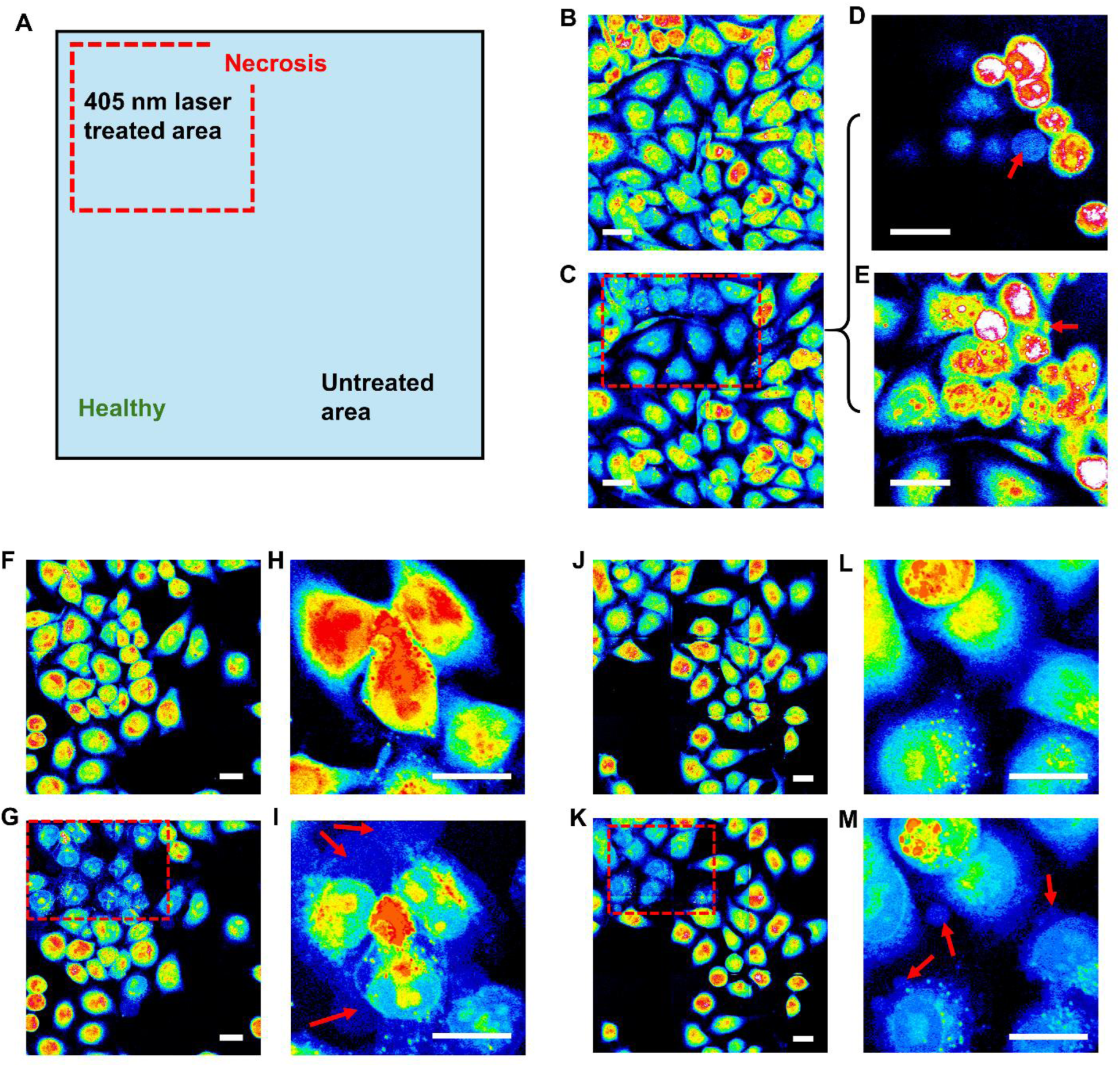
Induction of necrosis using 405 nm laser radiation. (A) Schematic illustrating regions of a culture dish following targeted illumination with a 405 nm continuous-wave laser. (B) SRS image of CHO-K1 cells prior to blue light treatment. (C) The same field of view after treatment with a 2 mW 405 nm laser for 80 seconds (50 frames). (D) Different focal planes of the 405 nm laser-treated region, highlighting necrotic blebs. (F, G) SRS images of HeLa cells before and after 405 nm laser treatment. (H, I) Three-dimensional projections of cells in the treated region before and after blue light exposure, showing prominent necrotic blebs post-treatment. (J-M) Similar analyses to panels F-I, performed with a lower laser dosage using 1 mW 405 nm laser power. Scale bars, 20 µm in all panels.

The same blue light treatment was applied to HeLa cells (transfected with enhanced green fluorescence protein conjugated with end-binding protein 3, EB3-EGFP), which exhibited a more pronounced decrease in SRS intensity compared to CHO cells, as shown in **Figures 5F–I**. HeLa cells also displayed significantly larger membrane blebbing and swelling under the same treatment conditions (**Figures 5F-I, Supporting Videos 1,2**). At lower 405 nm laser intensities, necrotic transitions in HeLa cells showed smaller blebs but remained more pronounced than those observed in CHO cells.

The reduction in SRS signals suggests that during necrosis, cellular proteins are released into blebs due to membrane damage. The weaker SRS signals in these blebs and the cytoplasm indicate water influx into the blebs and cells, diluting protein concentrations in these regions. This behavior contrasts sharply with apoptosis, where protein concentrations increase due to cytoplasmic condensation. These distinct transitions highlight the fundamental differences between necrotic and apoptotic processes.

In addition to monitoring intensity changes, SRS microscopy enables the observation of intracellular organelle dynamics, such as LD mobility. This analysis has been detailed in published studies.^29, 48^ While LD mobility analysis does not provide single-cell-level information, it can serve as a complementary approach to confirm transitions from healthy to apoptotic and necrotic states. Healthy cells are characterized by highly dynamic LDs, apoptotic cells exhibit significantly reduced LD mobility, and necrotic cells completely lose LD mobility (**Figure S7**).

### Analysis of apoptosis progression in suspended CHO cells

In biopharmaceutical production of recombinant proteins, CHO cells are cultured in bioreactors in a suspended state. To evaluate whether our apoptosis analysis based on SRS microscopy is applicable to suspended cells, we treated suspended CHO cells with staurosporine and analyzed their intensity and size changes as a function of treatment time. Similar to adherent cells, we observed an increase in average SRS intensity (**Figures 6A, B**) and a reduction in cell size (**Figure 6C**). While these changes were less pronounced compared to adherent cells, they were still clearly distinguishable. Notably, the changes did not exhibit a continuous increase over the treatment period, likely reflecting differences in cellular responses between the adherent and suspended states. The CHO-K1 cells used in this study are typically grown in 2D culture adhering to dish surfaces. To obtain suspended cells, we applied trypsin to detach them from the substrate. The transition to suspension may introduce additional stress, resulting in distinct responses compared to adherent cells. Currently, we lack the resources to culture suspended CHO cells under conditions that fully mimic bioreactor environments in the lab. Nevertheless, our findings demonstrate that apoptotic transitions in suspended cells can be detected and exhibit similar patterns to those observed in adherent cells.

**Figure 6.**
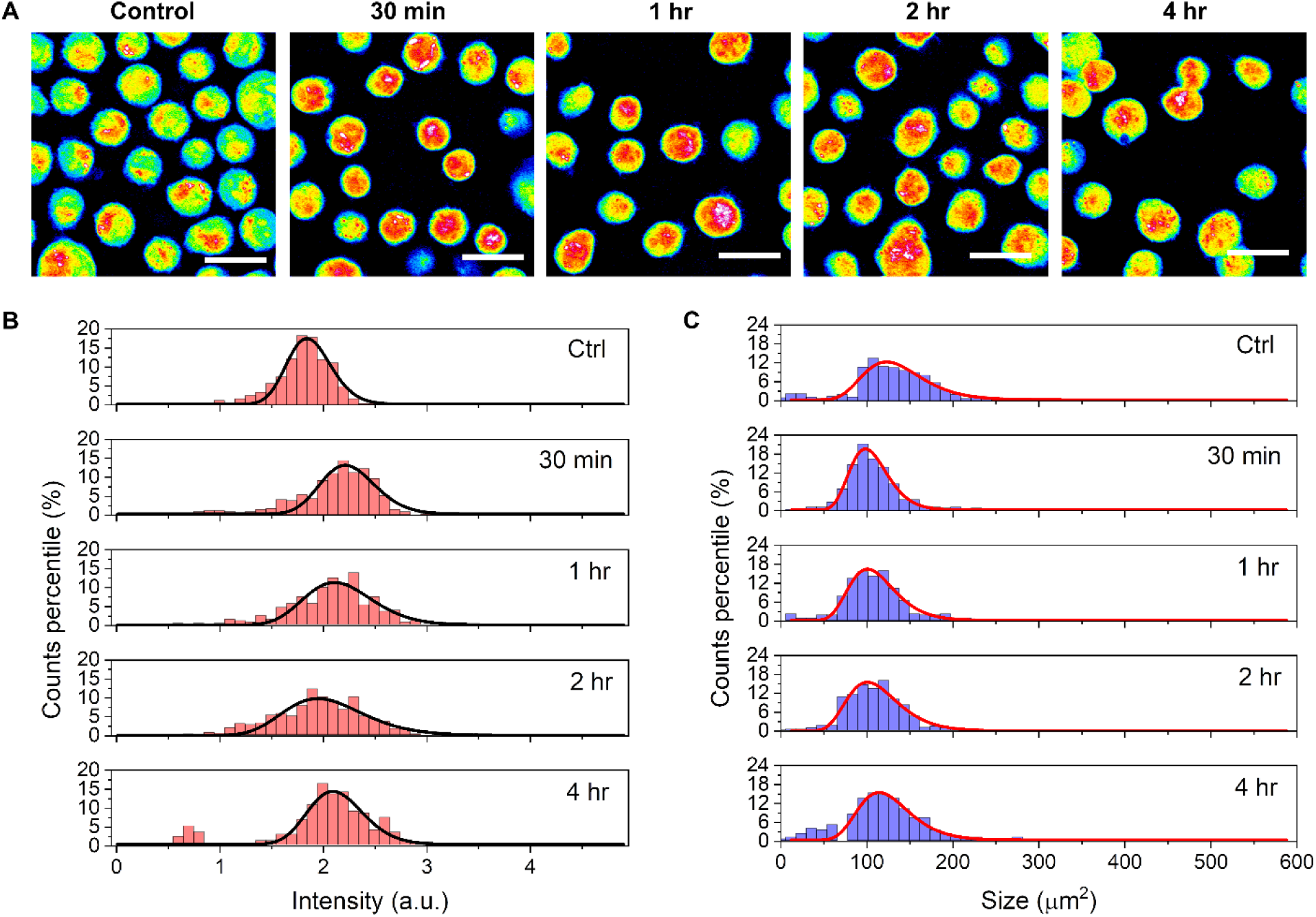
SRS detects increased protein concentration during apoptosis of suspended CHO cells. (A) SRS images of suspended CHO-K1 cells under untreated (control) and 1 µM staurosporine-treated conditions, captured at time points ranging from 30 minutes to 4 hours. Images are displayed using a jet color scheme. (B) Histograms depicting the distribution of cellular average SRS intensity, with lognormal function fits. (C) Histograms showing the distribution of cell sizes, also fitted with lognormal functions. Scale bars, 20 µm in panel A.

### Conclusions

We found that SRS microscopy combined with single-cell analysis enables the distinction between apoptotic and necrotic transitions, which exhibit distinct phenotypic and chemotypic changes. These differences are characterized by variations in intracellular protein concentration, cell size, and shape during different cell death processes. A schematic representation of these changes is shown in **Figure 7A**. In general, both apoptosis and necrosis involve a reduction in cell size. However, apoptosis is typically marked by an increase in overall protein concentration due to water loss, while necrosis shows a decrease in protein concentration caused by protein leakage and water influx as a result of membrane damage. These differences enable the detection of apoptotic and necrotic transitions using a 2D intensity versus cell size plot derived from SRS imaging, as summarized in **Figure 7B**. Furthermore, apoptotic bodies generally exhibit average SRS intensities similar to those of healthy cells, while necrotic blebs display weaker protein signals due to compromised membrane integrity. AVD is likely driven by the activation of ion channels in the cell membrane, leading to ion efflux that disrupts osmotic balance and causes water to exit the cell.^49–50^ This process is typically induced by the activation of the caspase cascade by staurosporine or other stressors.^51–53^ In contrast, necrotic changes are characterized by membrane damage that permits free diffusion of water into the cell.^54–55^ These changes can be analyzed using biological assays such as the Lactate Dehydrogenase (LDH) release assay, Western blot analysis, and ELISA kits. Compared to these methods, our SRS microscopy approach enables the study of live-cell transitions without requiring complex sample pre-preparation, while preserving rich chemical information.

**Figure 7.**
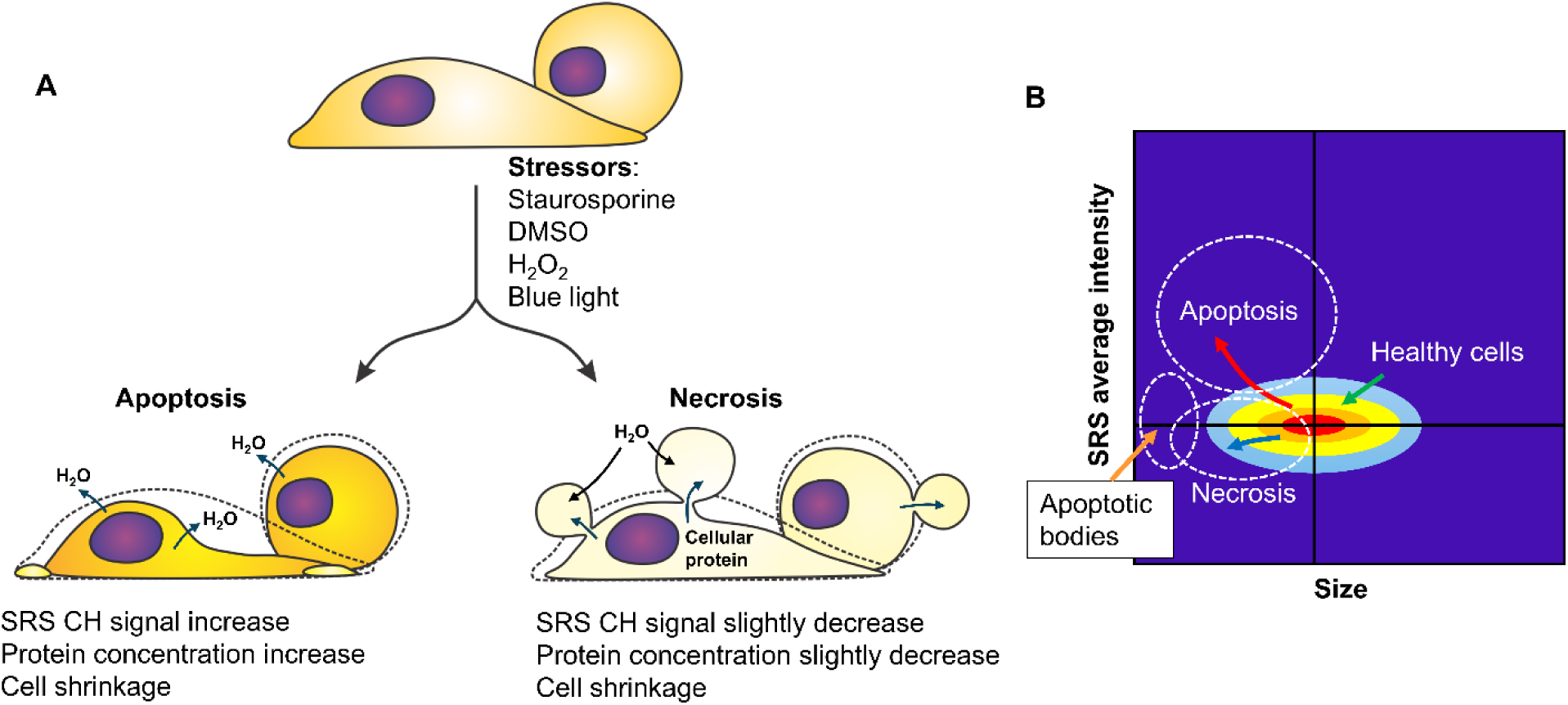
Detecting cell death pathways via SRS microscopy and single-cell quantitative analysis. (A) Schematic representation of phenotypic and chemotypic changes in cells undergoing apoptosis and necrosis. (B) Illustration of apoptosis- and necrosis-induced changes visualized on a 2D map of cell size versus average SRS intensity.

The single-cell analysis pipeline developed in this work is broadly applicable to studying cellular responses to various stressors or stimuli. Additionally, our findings highlight the potential of SRS microscopy as a label-free technology for evaluating cell death and responses to diverse perturbations. The chemically rich SRS signals provide critical single-cell-level information, allowing for the differentiation of subpopulations within mixed states, such as healthy, necrotic, and apoptotic cells. This method, when integrated with microfluidic platforms, is expected to facilitate rapid assessments of cell status and their correlation with CHO cell protein production in inline and online scenarios.

## Acknowledgment

This work is majorly supported by Merck Sharp & Dohme LLC, a subsidiary of Merck & Co., Inc., Rahway, NJ, USA, through a grant managed by the Merck-Purdue Center for Measurement Science granted to C.Z. It is also partially supported by NIH R35GM147092.

## Author Contributions

C.Z. Sh.M., B.W., and J.L. designed the project. Sh.M. and C.Z. performed the experiments. S.S., Sh.M., H.C., and C.Z. conducted the data analysis. Sh.M., K.M., J.L., B.W., and C.Z. attended biweekly meetings. K.M. helped with SRS system alignment and maintenance. Se.M. assisted in sample preparation and data collection. B.D. developed and tested the 3D imaging software for image acquisition and helped in imaging. C.Z. obtained the funding for this research.

