## Supplementary notes and figures for "Label-Free Quantification of Apoptosis and Necrosis Using Stimulated Raman Scattering Microscopy"

[3]Merck & Co. Inc.

### Quantitative single-cell analysis

#### Preprocessing and image analysis using ImageJ

To quantify the average SRS C-H signals of cells in each condition using ImageJ (as shown in Figure 1E), the 32-bit raw SRS image was first imported into ImageJ, and the intensity thresholding function was applied. Next, the "Analyze Particles" function was used with the particle size set to 100–infinity. The minimum size threshold of 100 pixels excluded random high-intensity specks and debris in the image that typically occupy only a few pixels. The average SRS intensity and standard deviation were calculated from 25 images per condition, generating the data for Figure 1E.

To create a lipid droplet (LD) mask from each image, the original C-H SRS image was subtracted from its Gaussian-blurred version (radius = 2 pixels, processed by ImageJ) using the “Image calculator function”. An intensity threshold was then applied to the resulting image followed by applying the "Analyze Particles" function to generate LD masks scaled from 0–255. These masks were divided by 255 using the “Math” function to create a normalized mask scaled from 0–1 (no mask area 0, mask area 1). The mask was multiplied by the original image to generate an LD-only image. Subtracting this LD-only image from the original C-H SRS image yielded LD-excluded SRS images, which primarily contained intracellular protein signals from cells, as shown in Figure 1C.

#### Cellpose segmentation of cells

The raw C-H SRS images were directly used as input for Cellpose 2.0. During processing, the average cell diameter was adjusted, and different Cellpose models were tested to ensure accurate identification of regions of interest (ROIs). The process was supervised, with manual intervention used to correct misidentified cells or include unassigned cells. The resulting outlines were saved as a .zip archive of ROI files, and the generated mask was saved as a PNG file.

The .zip mask could be imported into the ImageJ ROI Manager. By creating a blank image with the same dimensions as the original, the “Show All” function in the ROI Manager was applied, followed by the “Draw” function to create a cell mask scaled from 0–255. This mask was then normalized by dividing by 255 using the “Math” function and multiplied by the original image to generate a cell-only image. The resulting mask was directly used in CellProfiler for single-cell analysis.

#### CellProfiler-based single-cell analysis

It is important to note that accurate cell segmentation cannot be achieved using CellProfiler alone due to the absence of nucleus labeling in the cells.

For single-cell analysis, the cell mask obtained from Cellpose and the original C-H SRS image were used as inputs for CellProfiler. Additionally, the LD image or the LD-excluded image could be included as input. These images were appropriately named to allow CellProfiler to automatically select them based on their filenames.

The CellProfiler pipeline included the following modules: Identify Primary Objects, Measure Object Size and Shape, Measure Area Occupied, Measure Object Intensity, and Export to Spreadsheet. The typical diameter of primary objects was set to 10–80 pixels, and the manual threshold was set to 0.5. The smoothing scale was set to 0, the method for distinguishing clumped objects was set to "shape," and the method for drawing dividing lines between clumped objects was also set to "shape." The "Suppress local maximum distance" parameter was set to 30.

Parameters required for this study were selected using the "Select Measurements" function in the Export to Spreadsheet tab. The output Excel file from CellProfiler contained key information about the primary objects identified in the images. The total LD content could be calculated by using the LD image in conjunction with the cell mask input.

#### Generation of histograms and 2D density contour plots

The parameters of interest were selected from the output table saved in the Excel spreadsheet. All histograms were plotted with 50 bins. For intensity plots, the scale was set from 0 to 5. For size plots, the scale was set from 0 to 15,000 pixels, corresponding to 0–600 $\mu m^2$. For eccentricity, the scale ranged from 0 to 1, where lower values indicated shapes more similar to a circle. For total intensity plots, the scale was set from 0 to 50,000.

The plotted histograms are normalized by dividing the 'counts' in each bin by the total number of counts and multiplying by 100. This ensures that the total area of the histogram sums to 100%. Each bar represents the percentage of cells corresponding to the designated value on the x-axis.

The histograms were fitted using single, dual, or tri-lognormal functions. The functional forms were as follows:

y=y0 + A1/(sqrt (2*pi)*w1*x)exp(-(ln(x/xc1))^2/(2*w1^2))

y=y0 + A1/(sqrt (2*pi)*w1*x)exp(-(ln(x/xc1))^2/(2*w1^2))

+ A2/(sqrt (2*pi)*w2*x)exp(-(ln(x/xc2))^2/(2*w2^2))

y=y0 + A1/(sqrt (2*pi)*w1*x)exp(-(ln(x/xc1))^2/(2*w1^2))

+ A2/(sqrt (2*pi)*w2*x)exp(-(ln(x/xc2))^2/(2*w2^2))

+ A3/(sqrt (2*pi)*w3*x)exp(-(ln(x/xc3))^2/(2*w3^2))

The fitting parameters include y0, A1 (A2, A3), xc1 (xc2, xc3), w1 (w2, w3). A represents the relative number of events and xc represents the median value of the distribution.

The fitting was performed using the “Nonlinear Curve Fit” function in OriginPro 2020.

2D density contour plots were generated by correlating columns of parameters in the Excel spreadsheet. The data were processed using custom MATLAB scripts. The plot ranges for the parameters were consistent with those used in the histograms. To enhance the visualization of population clusters, the plots were rendered using the Jet color scheme.

#### LD mobility analysis

The mobility of LDs was analyzed using 50-frame time-lapse C-H SRS images. The "Particle Tracker" plugin in ImageJ was employed to quantify and save all trajectories. Trajectories from the same condition but different image stacks were combined to generate histogram plots.

To use the "Particle Tracker" plugin, the images were rescaled to 8-bit. The parameters were set as follows: radius = 3, cutoff = 0, percentile = (1–3), Link Range = 1, and Displacement = 5. The maximum displacement of LDs was analyzed using custom MATLAB scripts. Trajectories spanning fewer than 15 frames were filtered out by the MATLAB code.

Histograms of maximum displacement were generated using ImageJ and Origin2020.

### An explanation of the total SRS intensity increase during apoptosis

The cell shrinkage occurs in 3D, and the SRS image quantifies cell size in 2D. Assuming the 2D size of a cell is A and the height of the cell is h. Then the density of the cell

$$\rho \propto \frac{m}{V} \propto \frac{m}{A \times h}$$

Here m is the total mass of the cell

The measured SRS intensity

$$I \propto A \times \rho \propto A \times \frac{m}{A \times h} = \frac{m}{h}$$

Assuming the total mass of the cell remains constant, a decrease in the cell's height h can lead to an increase in the total integrated intensity measured by SRS. This phenomenon is observed during the early stages of staurosporine treatment in CHO cells, as illustrated in Figure S3. However, as the total mass of the cell decreases over time, the integrated intensity eventually declines. During necrosis, the total intensity of the cell decreases significantly.

### Supporting Figures

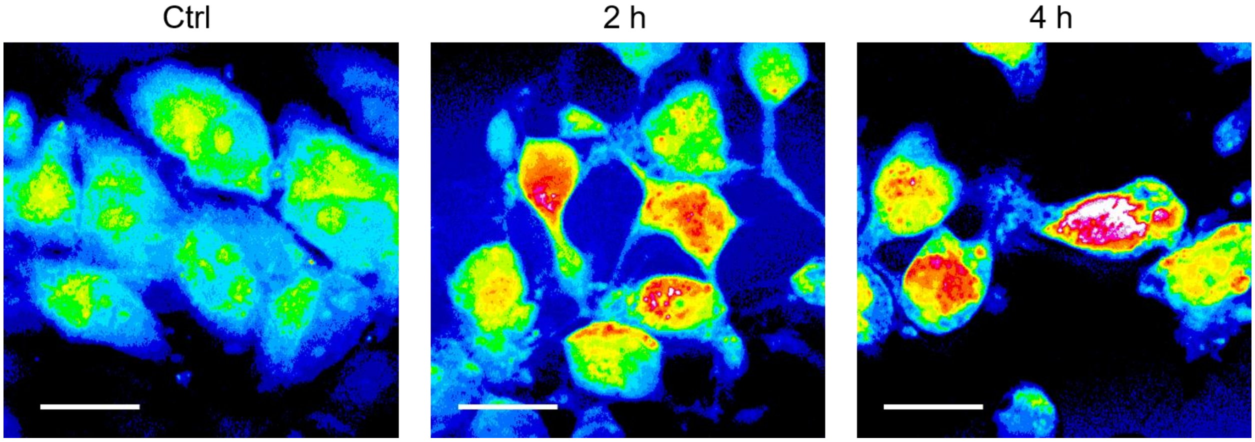


Figure S1. SRS intensity changes of HeLa cells in apoptosis. From left to right, untreated HeLa cells, HeLa cells treated with 1 μM staurosporine for 2 hours, HeLa cells treated with 1 μM staurosporine for 4 hours. Scale bars: 20 μm.

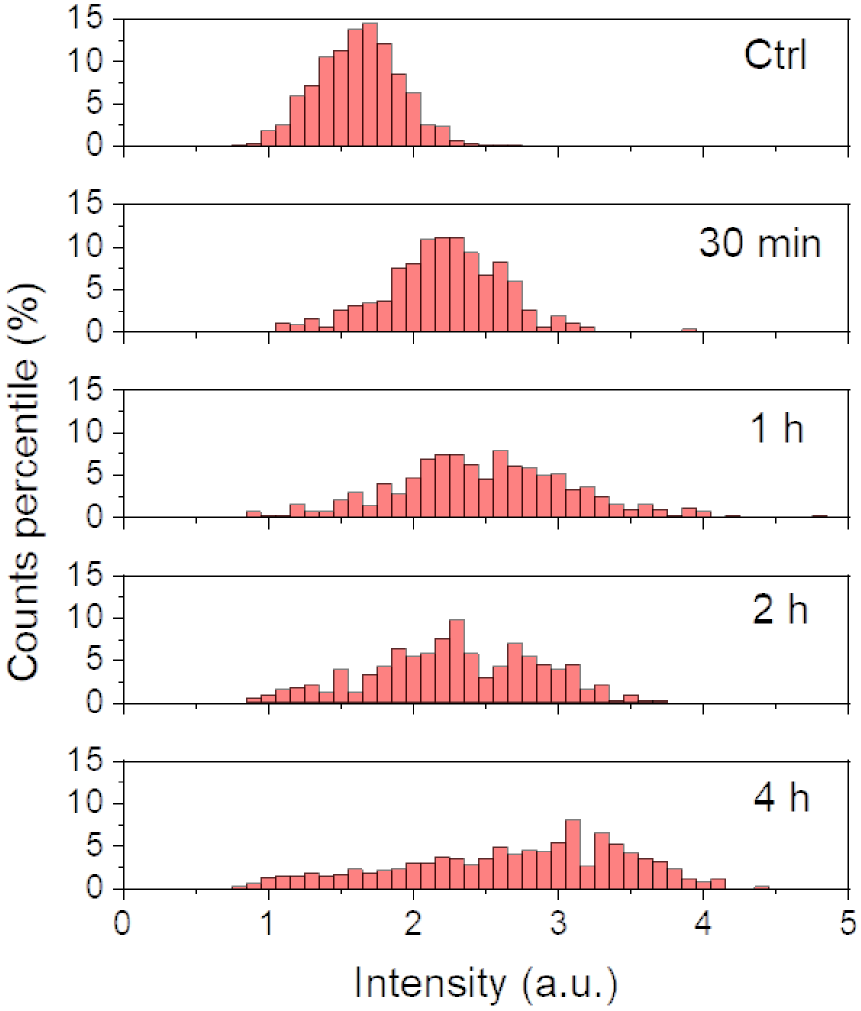


Figure S2. Histograms showing the distribution of cellular averaged SRS intensity after excluding lipid droplets from adherent CHO-K1 cells untreated and treated by 1 μM staurosporine for different time lengths.

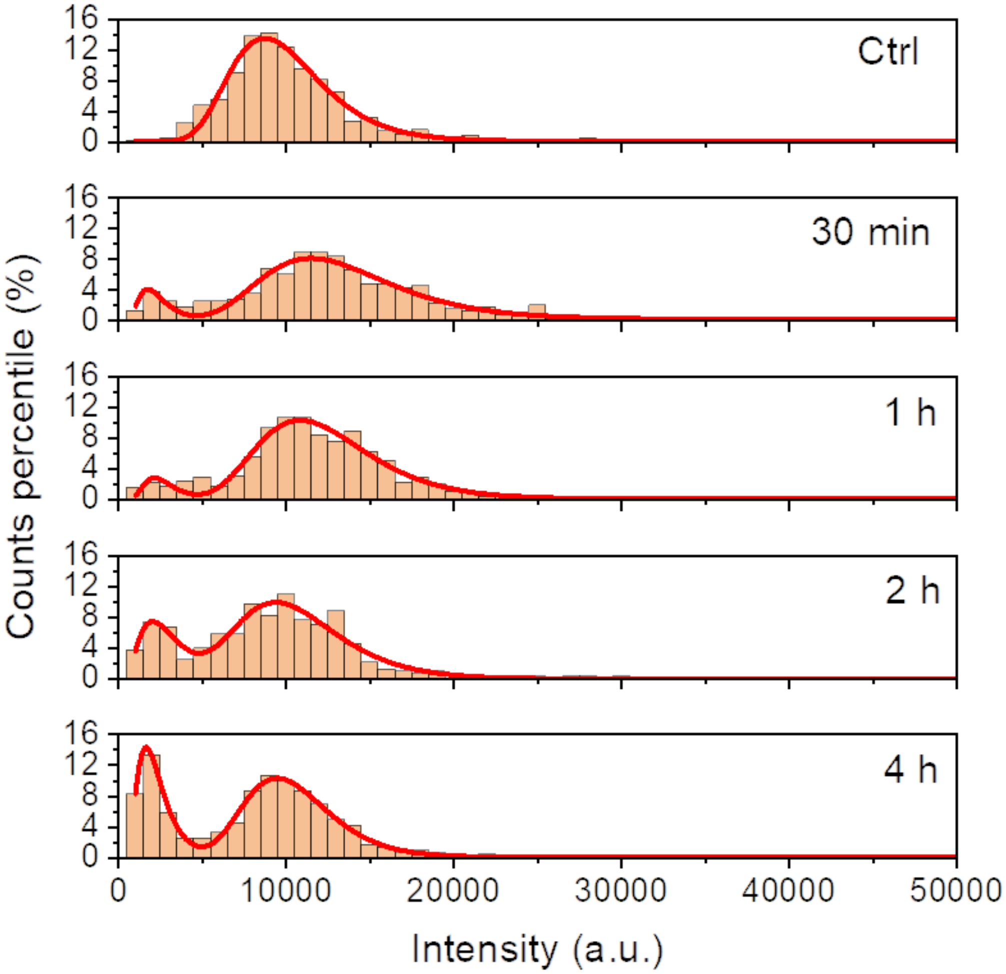


Figure S3. Histograms showing the distribution of integrated cellular SRS intensity for adherent CHO-K1 cells untreated and treated by 1 μM staurosporine for different time lengths. Red curves are fit using the lognormal function for the control (untreated group), and the dual-lognormal function for the rest.

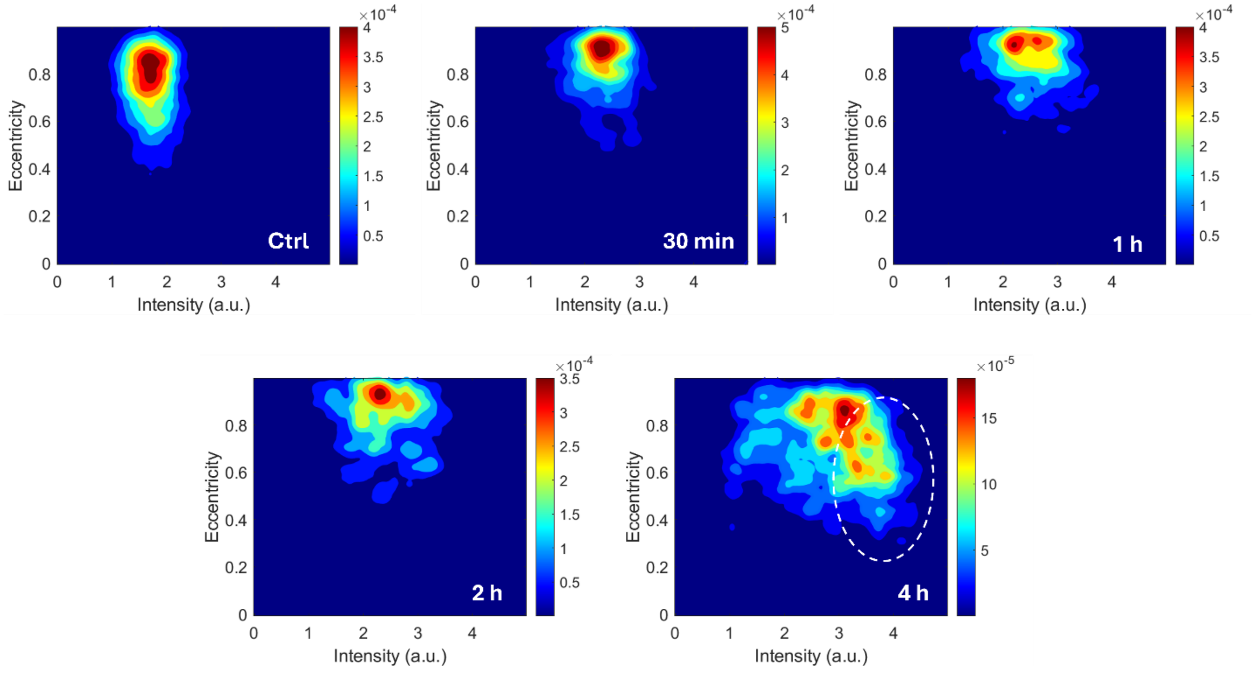


Figure S4. Two-dimensional contour plots depicting the relationship between averaged SRS intensity of cells and the eccentricity of cells untreated and treated with 1 μM staurosporine at different time points. The outlined area corresponds to more round-up late-stage apoptotic cells.

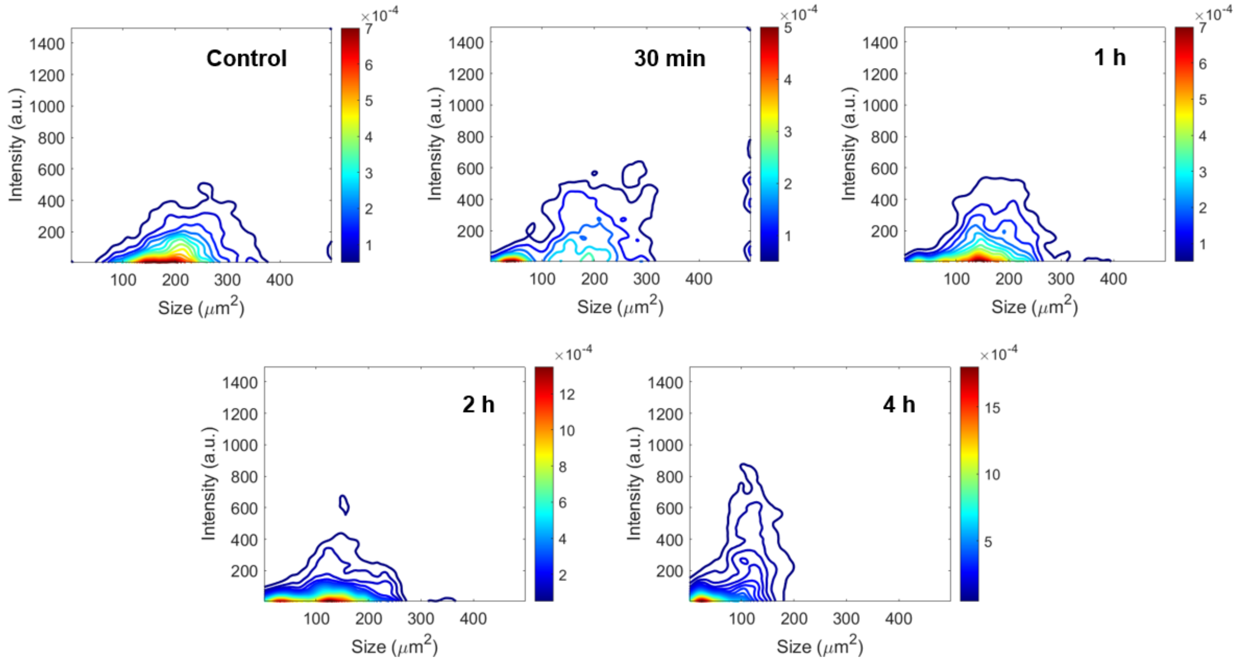


Figure S5. Two-dimensional contour plots depicting the relationship between SRS intensity from total lipid droplets in each cell and cell size under untreated and various apoptotic conditions induced by staurosporine.

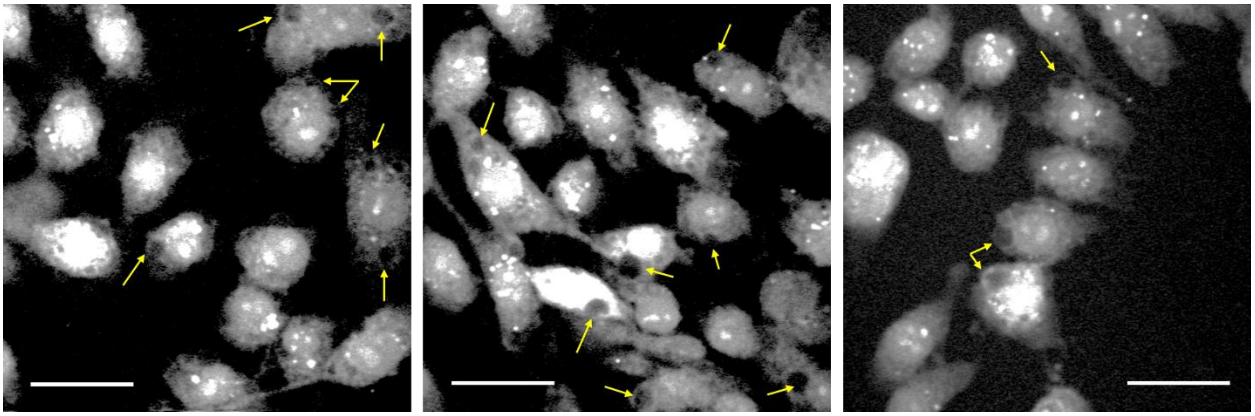

Figure S6. SRS images of CHO cells treated with 5 mM $H_2O_2$ for 4 hours. Arrows point out vacuoles and membrane blebs formed inside cells. Scale bars: 20 μm.

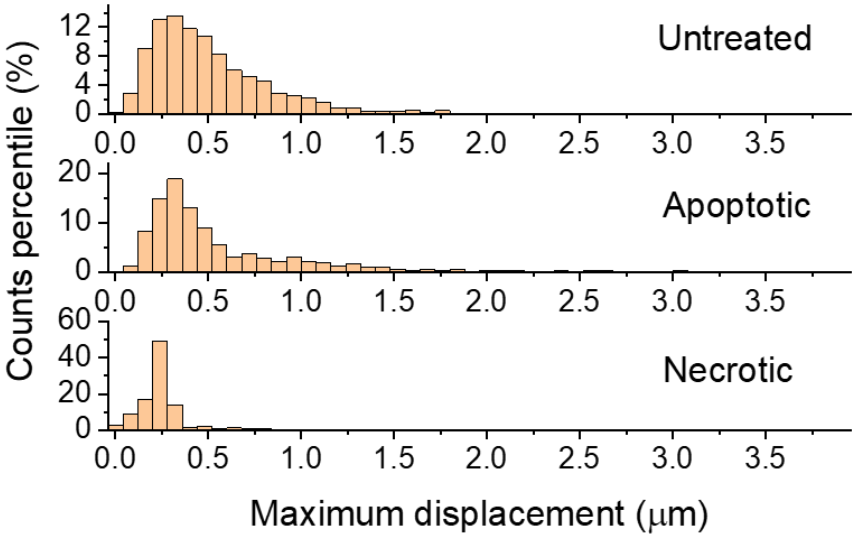


Figure S7. Histograms of lipid droplet maximum displacement of untreated, apoptotic, and necrotic cells. The apoptosis is induced by staurosporine and measured at 30 min to 1 hour after treatment. The necrosis is induced by high DMSO concentration and measured at 30 min to 1 hour after treatment.

Video S1. A 3D SRS image of HeLa cells before treatment with 405 nm laser.

Video S2. A 3D SRS image of HeLa cells after treatment with 2 mW 405 nm laser for 80 seconds.
